# CTFFIND4: Fast and accurate defocus estimation from electron micrographs

**DOI:** 10.1101/020917

**Authors:** Alexis Rohou, Nikolaus Grigorieff

## Abstract

CTFFIND is a widely-used program for the estimation of objective lens defocus parameters from transmission electron micrographs. Defocus parameters are estimated by fitting a model of the microscope’s contrast transfer function (CTF) to an image’s amplitude spectrum. Here we describe modifications to the algorithm which make it significantly faster and more suitable for use with images collected using modern technologies such as dose fractionation and phase plates. We show that this new version preserves the accuracy of the original algorithm while allowing for higher throughput. We also describe a measure of the quality of the fit as a function of spatial frequency and suggest this can be used to define the highest resolution at which CTF oscillations were successfully modeled.

## 1. Introduction

Estimating the electron microscope’s objective lens defocus and astigmatism from micrographs is of great importance in the three-dimensional reconstruction of biological specimens (reviewed by Cheng et al., 2015). Many programs are available for this purpose. Of those, CTFFIND3 (Mindell and Grigorieff, 2003) is widely used and thought to perform well under a range of circumstances (Marabini et al., 2015).

We present an updated version of this program, called CTFFIND4, which aims to match the quality of results obtained with CTFFIND3 and to do so significantly faster.

Like its predecessor, CTFFIND4 models the microscope’s contrast transfer function (CTF) as a 2-dimensional function of the spatial frequency vector **g**:

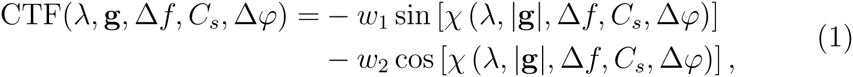

where the frequency-dependent phase shift *χ* is a function of the electron wavelength *λ*, the objective defocus Δ*f* and the spherical aberration *C*_*s*_,

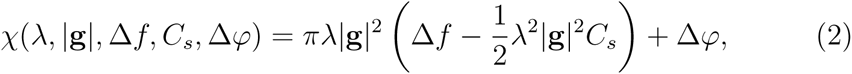

and Δ*φ* is an additional phase shift introduced by a phase plate, in the absence of which Δ*φ* = 0. *w*_2_ is the fraction of total contrast attributed to amplitude contrast, arising for example from electrons scattered outside the objective aperture or those removed by energy filtering (Yonekura et al., 2006). The value of this parameter depends on the specimen characteristics (e.g. ice thickness, heavy metal stains) as well as microscope properties (e.g. acceleration voltage, diameter of the objective apverture) and must be given by the user. The relative phase contrast is 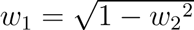.

Equation 1 can be simplified and made more computationally efficient (Fernando and Fuller, 2007):

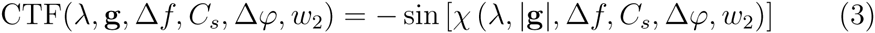

with

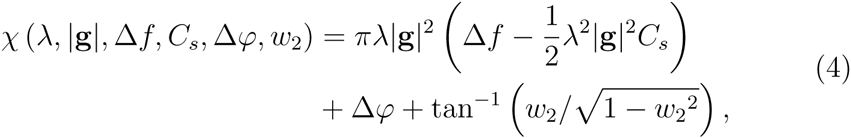

where the tan^-1^ term can be precomputed, so that evaluating the CTF only requires one trigonometric function call.

To account for astigmatism of the objective lens two defocus values, Δ*f*_1_ and Δ*f*_2_, are defined, which describe the lens’ defocus along normal directions:

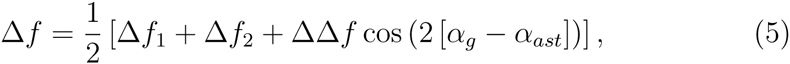

where *α*_*g*_ is the angle between **g** and the *X* axis in reciprocal space. The astigmatism is defined by its magnitude ΔΔ*f* = Δ*f*_1_ - Δ*f*_2_ and its polar angle *α*_*ast*_, the angle between the image *X* axis and the direction along which Δ*f* = Δ*f*_1_ (Figure 1).

**Figure 1:**
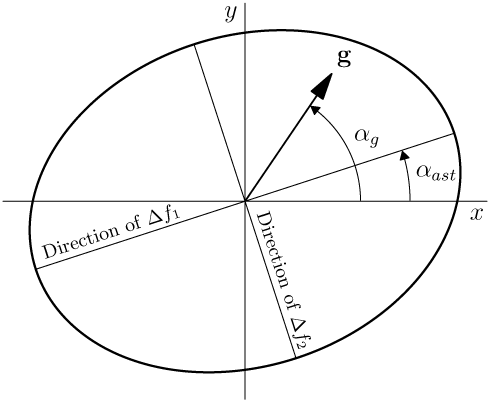
Two defocus values, Δ*f*1 and Δ*f*2, and an angle, *α*_*ast*_ define an astigmatic CTF. The effective defocus at an arbitrary point **g** (scattering vector) in reciprocal space is defined by Equation 5. Adapted from Figure 3 of Mindell and Grigorieff (2003).

## 2. User input

By default, input parameter values are given interactively, with a question-answer sequence inspired by that of the IMAGIC image processing package (van Heel et al., 1996), including help messages (given in response to “?”) and default answers which are updated with the user’s previous answers.

Alternatively, the user can give the 

~~~
--old-school-input
~~~

 command-line option, in which case CTFFIND4 will accept the same input as CTFFIND3 but use the CTFFIND4 algorithms described here. This feature is meant to facilitate the use of CTFFIND4 in the context of pre-existing scripts and workflows, though it does not allow access to new features such as movie processing or phase shift determination.

Aside from the obvious input parameters (micrograph file name, microscope acceleration voltage etc.), the user must choose a number of parameter values (listed in Table 1), for which suggested defaults are built in to the program. These default values should be taken as guides only, but may serve as good starting points when searching for optimal input parameters.

**Table 1:** Input parameters whose values must be chosen by the user. The four parameters below the dashed line are only required when a stack of frames is given as input (*N*_*frames*_; see Section 3.7) or when the user indicates that a phase plate with variable phase shift was used (Δ*φ*_*min*_, Δ*φ*_*max*_, Δ*φ*_*step*_; see Section 3.8).

| Symbol | Description (default value, unit) |
| --- | --- |
| $w_2$ | Relative amplitude contrast (0.07) |
| $N_d$ | Size of spectrum (512 pixels) |
| $g_{min}^{-1}$ | Min. resolution in target function (30 Å) |
| $g_{max}^{-1}$ | Max. resolution in target function (5 Å) |
| $\Delta f_{min}$ | Lower bound of initial defocus search (5000 Å) |
| $\Delta f_{max}$ | Upper bound of initial defocus search (50000 Å) |
| $\Delta f_{step}$ | Step size of initial search for defocus (500 Å) |
| $\Delta\Delta f_{res}$ | Astigmatism restraint (100 Å) |
| $N_{frames}$ | Number of frames to average (1) |
| $\Delta\varphi_{min}$ | Lower bound of initial phase shift search (0 rad) |
| $\Delta\varphi_{max}$ | Upper bound of initial phase shift search (3.15 rad) |
| $\Delta\varphi_{step}$ | Step size of initial phase shift search (0.01 rad) |

Optionally, the user may supply pre-computed amplitude spectra (using command-line option 

~~~
--amplitude-spectrum-input
~~~

), in which case amplitude spectrum calculation (Section 3.1) will be bypassed. Background subtraction (Section 3.2) can also be bypassed by giving 

~~~
--filtered-amplitude-spectrum-input
~~~

.

## 3. Algorithm

CTFFIND4 mostly re-implements CTFFIND3 with a few modifications. To summarize, the algorithm consists of computing an amplitude spectrum from the input micrograph, estimating the spectrum’s background, subtracting this from the original spectrum, and evaluating the similarity between theoretical two-dimensional CTF functions and the remaining oscillatory signal. The parameters for the theoretical CTF are varied until the similarity is maximized, yielding an estimate of the microscope’s defocus and astigmatism parameters.

### 3.1. Amplitude spectrum

The amplitude spectrum is computed by taking the absolute of the Fourier transform of the whole micrograph (padded to square dimensions if necessary). The amplitude spectrum is then down-sampled to the desired dimensions by Fourier truncation, which discards terms outside a user-defined central part of the Fourier transform of the amplitude spectrum.

The size of the decimated amplitude spectrum, *N*_*d*_, is chosen by the user. To lower the computational cost of the scoring function, it should be kept as small as possible (see section 3.3). However, a small spectrum may not describe CTF oscillations accurately enough, leading to increased errors in parameter estimates. This is epecially true in cases of large defocus (for a more quantitative description of this effect, see Penczek et al., 2014). The default spectrum size in CTFFIND4 is 512 × 512 pixels, which is sufficient in the majority of cases, but in many instances users may wish to use smaller dimensions (e.g. 256 × 256) for computational efficiency.

### 3.2. Background subtraction

Most of the power in experimental images is concentrated in the lowest spatial frequencies, giving a dominant peak centered at the origin of the amplitude spectrum and a slow-decreasing ramp towards high frequencies, on top of which amplitude oscillations due to the CTF are sometimes barely discernible. To estimate this ramp, both versions of CTFFIND use the same box-convolution algorithm: the down-sampled amplitude spectrum is convoluted in real space with a square boxcar function, and the resulting smooth spectrum is subtracted from the signal.

The convolution operation is carried out only at radii corresponding to frequencies greater than *g*_*min*_. Pixels near the origin are left unchanged, so that after subtraction of the smoothed spectrum they are set to 0. In addition, the central 3 pixels in each direction (*x* and *y*) are ignored by the convolution kernel, so that any artefacts in the central cross of the amplitude spectrum do not affect the quality of background subtraction.

### 3.3. Scoring function

We use the normalized cross correlation coefficient between the CTF and the experimental, decimated and background-subtracted amplitude spectrum **A_d_** as a target function for our search and refinement of CTF parameters:

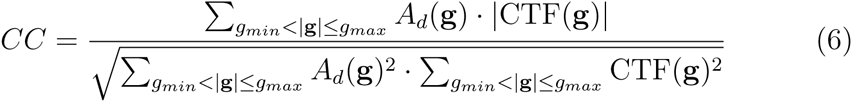

The cross-correlation is computed by iterating over all pixels in **A_d_** which lie between the radii corresponding to spatial frequencies *g*_*min*_ and *g*_*max*_. The computational cost of evaluating this function is therefore proportional to 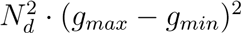.

In some cases CTF oscillations are not clearly detectable above background noise, such that cross-correlation values are low and they do not discriminate sufficiently between the correct (unknown) set of parameter values and alternatives. Because microscopists commonly aim to minimize astigmatism during data collection however, it is often helpful to prefer CTF parameter values which imply lower astigmatism. To this end, the user can place a restraint on ΔΔ*f*, the amplitude of astigmatism, by specifying ΔΔ*f*_*res*_. The final score *S* is then:

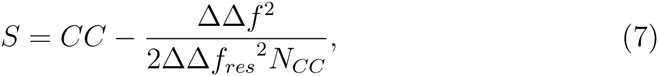

where *N*_*CC*_ is the number of pixels which were included in the computation of *CC*. The restraint (second) term in Equation 7 will penalize parameter values which imply very astigmatic CTFs and this penalty will be especially significant when CTF oscillations are weak so that CC tends to be small. The strength of the penalty will be greater with lower ΔΔ*f*_*res*_ and larger astigmatism ΔΔ*f*. In CTFFIND4, this restraint can be turned off by setting ΔΔ*f*_*res*_ *<* 0, in which case *S* = *CC*. If ΔΔ*f*_*res*_ is set to zero by the user, it is internally set to a small value (100 Å, corresponding to a very strong restraint) to avoid division by zero.

### 3.4. Search for astigmatism angle

During execution of CTFFIND3, most of the computation time is spent on an initial 3-dimensional exhaustive search over Δ*f*_1_, Δ*f*_2_ and *α*_*ast*_. To make our program more efficient, we first estimate *α*_*ast*_ separately, using an algorithm described by van Heel et al. (2000). First, the preprocessed amplitude spectrum **A_d_** is mirrored along one of its axes. It is then aligned rotationally against its mirrored self using a 1-dimensional exhaustive search with 5^°^ steps. *α*_*ast*_ is then taken to be half of the rotation angle which relates the two mirror images.

### 3.5. Search for defocus values

In the next step, *α*_*ast*_ is fixed and an exhaustive 2-dimensional search is done between Δ*f*_*min*_ and Δ*f*_*max*_ to find the values of Δ*f*_1_ and Δ*f*_2_ which maximize *S*.

### 3.6. Refinement of astigmatism & defocus value

The final step of the algorithm consists of a 3-dimensional conjugate-gradient maximization of *S*, which yields the final estimates of the values Δ*f*_1_, Δ*f*_2_ and *α*_*ast*_.

### 3.7. Processing dose-fractionated movies

Under some circumstances, improved CTF oscillations may be recovered from experimental micrographs by averaging amplitude spectra of movie frames rather than computing the amplitude spectrum from the sum of aligned frames (Bartesaghi et al., 2014). McMullan et al. (2015) observed a similar phenomenon in their study of Thon rings from amorphous ice and suggested an optimal dose of 4 e^-^/Å^2^ to observe oscillation around 3.7 Å.

CTFFIND4 supports CTF estimation from movies by allowing the user to give a stack of frames as input and specify *N*_*frames*_, the number of frames to average together before computing amplitude spectra. If the user sets *N*_*frames*_ = 1, amplitude spectra are computed from each frame and then averaged.

### 3.8. Processing micrographs recorded using phase plates

Volta phase plates (Danev et al., 2014) have recently become available commercially. Unlike other phase plate designs, they introduce a phase shift of scattered relative to unscattered electrons which is variable over time/irradiation and may therefore need to be measured during data collection and/or *a posteriori* from the collected images. We added a phase shift term Δ*φ* to our CTF model (see Equation 2), which can be fit to **A_d_** simultaneously with Δ*f*_1_ and Δ*f*_2_ so that users may estimate their phase plate’s phase shift. If the user specifies that a phase shift should be fit, *α*_*ast*_ is estimated first as usual (see Section 3.4), but the exhaustive search is then 3-dimensional (Δ*f*_1_, Δ*f*_2_, Δ*φ*) and the subsequent maximization is 4-dimensional (including *α*_*ast*_).

## 4. Outputs

A summary text file is output, to which the final estimates of Δ*f*_1_, Δ*f*_2_, *α*_*ast*_ and Δ*φ* for each input micrograph are written (one line per micrograph), as well as the final cross-correlation between **A_d_** and the fit CTF (Equation 7). The other outputs of CTFFIND4 are concerned with giving the user feedback regarding the quality of the fit.

### 4.1. Diagnostic image

A diagnostic image is generated, showing |**CTF_fit_**| overlaid onto a version of **A_d_** thresholded to improve contrast of the Thon rings (Figure 2). This is meant to provide qualitative feedback and relies on the user’s expertise to judge whether the fit was satisfactory and/or until which resolution the fit was successful.

**Figure 2:**
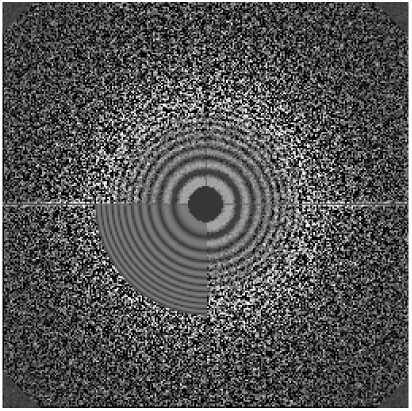
Diagnostic image from micrograph #1 of set #7 of the CTF challenge, output by CTFFIND4 using runtime parameters detailed in Table 3. The 2-dimensional CTF (**CTF_fit_**) is overlayed onto the preprocessed amplitude spectrum (**A_d_**) up to the radius corresponding to *g*_*max*_.

### 4.2. Diagnostic fit profile

CTFFIND4 also computes 1D profiles of **A_d_** and |**CTF_fit_**|, **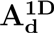** and |**CTF_fit_**|**^1D^**. In the simple case where ΔΔ*f* = 0, these are simply radial averages with bin width *b*, e.g.:

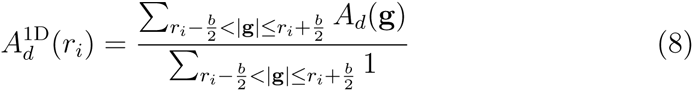

Evaluating these 1D profiles in the case of astigmatic CTF functions (ΔΔ*f* ≠ 0) is not trivial and this problem has been approached by several authors (e.g. Mallick et al., 2005). Our approach’s first step is to count, for every pixel, the number *n* of CTF extrema between it and the origin of the amplitude spectrum image. To find the spatial frequencies of CTF extrema requires solving the derivative of Equation 3 with respect to *χ*,

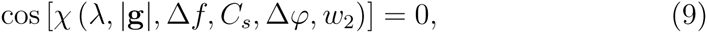

for which there is an infinite number of solution, exactly *n* of which are below any given spatial frequency |**g**|:

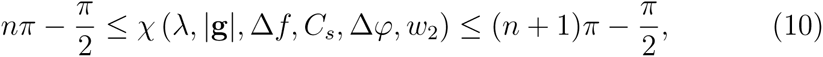

so that

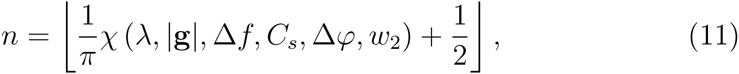

where ⌊⌋ denotes the floor function. We hold this count in memory as image **E** (Figure 3).

**Figure 3:**
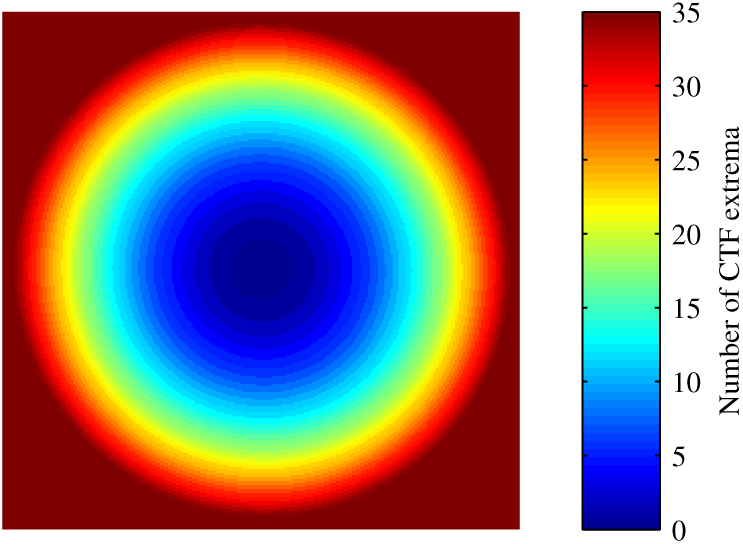
Image **E** generated from the CTF fit to micrograph #1 of set #7 of the CTF challenge. At every pixel (corresponding to a spatial frequency vector **g**), this image records *n*, the number of preceding CTF extrema (Equation 11). Here this value is color-coded, so that pixels at spatial frequencies before the first extremum of the CTF, which have value 0, are displayed in dark blue. Pixels that have 35 or more preceding CTF extrema are shown in dark red.

This allows the grouping of pixels according to their “position” along the CTF (number of preceding extrema), as opposed to their spatial frequency (distance from the origin)^1^ and thus average them regardless of astigmatism. Specifically, we wish to compute the 1D profile of **A_d_** along the direction of average defocus *α*_*mid*_ = *α*_*ast*_ + *π/*4, in bins of width *b* = 1 pixel centered at frequencies *r*_*i*_ indexed by 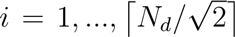. To this end we define *S*(*r*_*i*_), the set of **g** vectors which index pixels that can be averaged together into 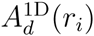 because they have the same number of preceding extrema [*E*(**g**)] and because their assigned CTF_fit_ values are closer to the value of CTF_fit_ along *α*_*mid*_ at *r*_*i*_ than at any other bin center *r*_*j*_ (*j ≠ i*):

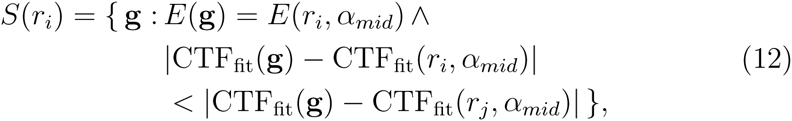

where we use set-builder notation; {*x*: *p*(*x*)} denotes the set of values of variable *x* such that *p*(*x*) is satisfied, ∧ denotes conjunction. The values of **A_d_** at pixels within set *S*(*r*_*i*_) can then be averaged to give 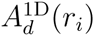:

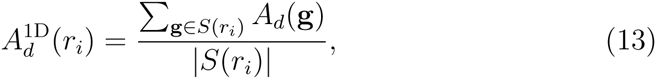

where *||* denotes the cardinality of a set. Similarly for |**CTF_fit_**|:

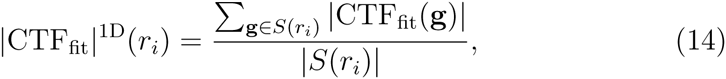

### 4.3. Estimating the quality of fit

In an attempt to provide a quantitative measure of the quality of fit, we implemented a spatial frequency-dependent measure of the correlation between |**CTF_fit_**|**^1D^** and 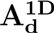, similar to that described by Huang et al. (2003). We chose the normalized cross-correlation, computed at intervals delimited by the maxima of |**CTF_fit_**|^**1D**^ along *α*_*mid*_:

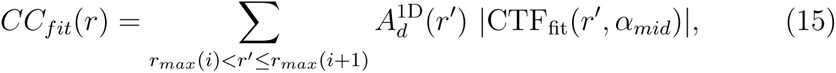

where we have omitted the usual normalization terms for clarity, and where *r*_*max*_(*i*) and *r*_*max*_(*i* + 1) are the frequencies of the CTF extrema immediately preceding and following *r*.

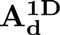, **CTF_fit_** and **CC_fit_** are output by CTFFIND4 as a text file which can be plotted using an accompanying script (Figure 4). In addition, an estimate is made of the highest resolution up to which a “good” fit to **A_d_** is obtained. We chose the criterion of *CC*_*fit*_(*r*) ≥ 0.75 heuristically for this purpose, with the hope that a criterion based on a statistical significance test may be derived and implemented in future versions of the software.

**Figure 4:**
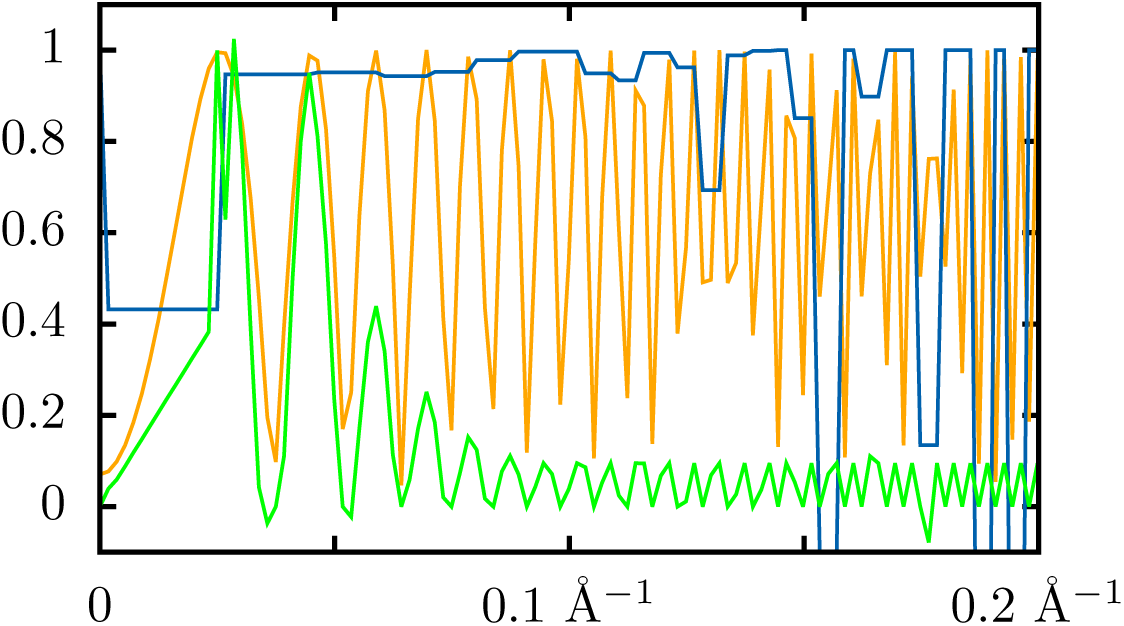
Output diagnostic plots describing the experimental amplitudes (**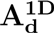**, green), the fit CTF (|**CTF_fit_**|^**1D**^, orange) and goodness of fit (**CC_fit_**, blue) for micrograph #1 of set #7 of the CTF challenge. For this micrograph, the final estimates were Δ*f*1 = 29070 Å, Δ*f*2 = 28313 Å and *α*_*ast*_ = 56.5^°^. The highest resolution at which Thon rings were deemed to be modeled correctly was 6.5 Å. The experimental amplitude profile (green) is normalized such that: the minima of the oscillations are set to 0.0; the second peak of the power spectrum (in this case at around 0.04) is 0.95; the maxima of oscillations are further normalized to 0.1 if their maxima would be *<*0.1 otherwise. Because of aliasing, one does not observe zeroes in |**CTF_fit_**|^**1D**^. One would normally solve this by increasing *N*_*d*_, but we restricted ourselves to previously-used parameter values for this experiment (see caption to Table 3 for more details).

## 5. Benchmarking

In the following paragraphs, CTFFIND versions 3.5 and 4.0.16 were used. Executable binaries and source code for both are available at http://grigoriefflab.janelia.org/ctf.

### 5.1. Speed

We expect that the time spent by CTFFIND4 evaluating *S*, the objective function, is proportional to 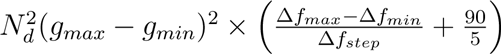 whereas for CTFFIND3 it is proportional to 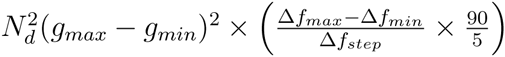 (note the multiplication in the last set of parentheses). The speedup in that part of the program should therefore be on the order of 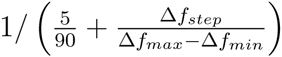. Other parts of the algorithm (such as the computation of **A_d_**) may also affect the speed-up somewhat.

We measured execution times of CTFFIND3 and CTFFIND4 with micro-graphs of bacteriorhodopsin, which had been used in benchmarking CTFFIND3 (Mindell and Grigorieff, 2003), as inputs. Using the same parameters as had been used by Mindell and Grigorieff, one would expect a 9-fold speedup in the search over *S*, but we only observed a *∼*3.7-fold speedup (Table 2). We assume that this is because under those circumstances, CTFFIND4 spends proportionally less time evaluating *S* and more time on other parts of the algorithm.

**Table 2:** Comparison of defocus parameter estimates from CTFFIND3 and CTFFIND4 to those obtained by crystallographic refinement of bacteriorhodopsin (Mindell and Grigorieff, 2003). The differences *E* between crystallographic values and those obtained by CTFFIND are given in nm (Δ*f*1, Δ*f*2) and degrees (*α*_*ast*_). Runtimes are given in seconds and were measured on a single Intel(r) Xeon(r) E5-287W CPU core operating at 3.10 GHz. To make runtimes comparable, the computation of 1D profiles and extra statistics were turned off in CTFFIND4 (they added *∼*2 seconds to the runtime). Speedups report the ratio of CTFFIND3 to CTFFIND4 runtimes. The following parameters were used: *N*_*d*_ = 128, 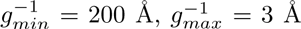, Δ*f*_*min*_ = 1000 Å, Δ*f*_*max*_ = 10000 Å, Δ*f*_*step*_ = 500 Å. The astigmatism restraint was turned off. The amplitude of astigmatism (ΔΔ*f*) was 727, 845, 884 and 812 Å, or 12%, 15%, 15% and 20% of Δ*f*1 for the four micrographs respectively. Profiling indicated that CTFFIND4 spent only *∼*15% of execution time evaluating *S*, which may explain why the measured speedup did not match the predicted 9-fold speedup.

| Micrograph | CTFFIND3 |  |  |  | CTFFIND4 |  |  |  |  | Speedup |
| --- | --- | --- | --- | --- | --- | --- | --- | --- | --- | --- |
| | $\epsilon(\Delta f_1)$ | $\epsilon(\Delta f_2)$ | $\epsilon(\alpha_{ast})$ | Runtime | $\epsilon(\Delta f_1)$ | $\epsilon(\Delta f_2)$ | $\epsilon(\alpha_{ast})$ | Runtime | $CC_{fit} = 0.75$ (Å) | |
| b3730 | 0.6 | -6.1 | -3.5 | 3.9 | 2.4 | -0.7 | -8.1 | 1.1 | 4.5 | 3.7 |
| b3736 | 10.0 | 1.8 | -3.5 | 3.9 | 12.7 | 2.0 | -4.0 | 1.1 | 3.2 | 3.7 |
| b3737 | -22.2 | 10.2 | -1.9 | 3.9 | 9.8 | 3.1 | -0.1 | 1.1 | 3.5 | 3.6 |
| b3739 | -0.3 | 0.9 | 1.1 | 3.9 | -0.6 | 2.0 | 1.2 | 1.1 | 3.1 | 3.6 |

As a separate test we ran both versions on a set of 24 micrographs of 3712 *×* 3712 pixels (set #3 from the CTF challenge, see below) using the default parameter values listed in Table 1 and 16 CPU threads. Under those circumstances, CTFFIND4 achieved speed-ups of ~10-fold relative to CTFFIND3 (*μ* = 9.85, *σ* = 0.24).

### 5.2. Accuracy

The micrographs of bacteriorhodopsin provide a useful benchmark for the accuracy of defocus parameter estimation because an earlier crystallographic study estimated their defocus parameters. We found that both versions of CTFFIND were similarly accurate, with errors on the order of a few nanometers (Table 2).

Assuming that CTFFIND3 provides accurate estimates of defocus parameters under most circumstances, we also aimed to ensure that CTFFIND4’s estimates closely matched those from CTFFIND3 under a wide range of circumstances, despite the significant algorithmic changes between the two versions. To this end, we ran both versions on all 9 sets of micrographs made available by Marabini et al. (2015) as part of the CTF challenge, and computed the percentage difference in estimates of defocus parameters between the two versions. We found discrepancies in defocus estimates were usually below 1%, but that discrepancies in *α*_*ast*_ were often large, in the tens of degrees (Table 3), presumably because under experimental conditions (high noise) and with low levels of astigmatism (ΔΔ*f /*Δ*f*_1_ on the order of 2%), this parameter is poorly determined by the Thon rings. Discrepancies in *α*_*ast*_ estimates were notably reduced for one of the sets of micrographs (#9), which consisted of simulated images and had the largest mean astigmatism. They were also very low (1.75 degrees on average) when processing bacteriorhodopsin micrographs, which have relatively large amounts of astigmatism (12 to 20% of Δ*f*_1_; Table 2).

In a significant departure from earlier versions, CTFFIND4 does not discard any parts of the input image when computing its amplitude spectrum. We reasoned that this feature, intended mainly to avoid artefacts in the Fourier transform due to sharp features such as photographic film labels or pieces of dust, was no longer necessary, since the vast majority of datasets are no longer recorded on film, and since features such as film labels can be removed by cropping the micrographs with any of the common image processing packages. In fact, we had noticed that when no features such as film labels were present in the input image, and when only some regions of the micrograph contained carbon film, CTFFIND3’s algoritm would often discard those regions. This reduced the amplitude of Thon rings and we hypothesized it might also reduce the accuracy of defocus parameter estimates, though we did not test this. In any case we did not see a need to maintain such a feature in new versions of CTFFIND.

**Table 3:**
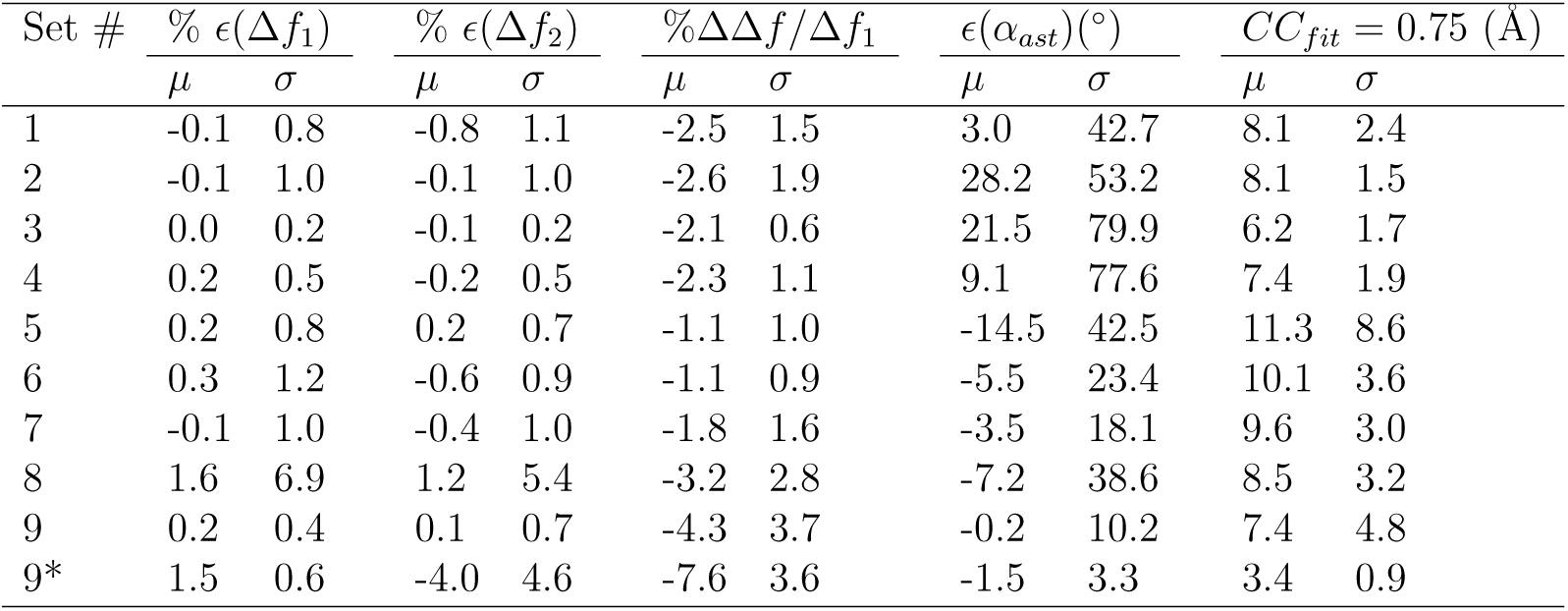
CTFFIND3 and CTFFIND4 find very similar defocus parameter values for CTF challenge micrographs. Nine sets of micrographs (described in Marabini et al., 2015) were processed with CTFFIND4, using the same parameters as had been used in one of the authors’ (N.G.) submission to the CTF challenge using CTFFIND3. Those parameters were: *N*_*d*_ = 256, 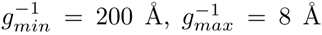, Δ*f*_*min*_ = 1000 Å, Δ*f*_*max*_ = 90000 Å, Δ*f*_*step*_ = 200 Å, ΔΔ*f*_*res*_ = 200 Å. For set #9, CTF parameters were also estimated with adjusted parameters *N*_*d*_ = 512 and ΔΔ*f*_*res*_ = 100 Å (which inactivates the astigmatism restraint) and the results are shown in line 9*. For each set, the mean (*μ*) and standard deviation (*σ*) of the difference *E* between CTFFIND 4 and CTFFIND3’s estimates of each of the defocus parameters (Δ*f*1, Δ*f*2, *α_ast_*) are shown as percentages or degrees, as are measures of the amount of astigmatism (ΔΔ*f /*Δ*f*1) and estimates of the highest resolution (in Å) at which Thon rings are reliably fit, as given by the *CC*_*fit*_ = 0.75 criterion.

| Set # | % $\epsilon(\Delta f_1)$ | | % $\epsilon(\Delta f_2)$ | | % $\Delta\Delta f/\Delta f_1$ | | $\epsilon(\alpha_{ast})(^\circ)$ | | $CC_{fit} = 0.75$ (Å) | |
| --- | --- | --- | --- | --- | --- | --- | --- | --- | --- | --- |
| | $\mu$ | $\sigma$ | $\mu$ | $\sigma$ | $\mu$ | $\sigma$ | $\mu$ | $\sigma$ | $\mu$ | $\sigma$ |
| 1 | -0.1 | 0.8 | -0.8 | 1.1 | -2.5 | 1.5 | 3.0 | 42.7 | 8.1 | 2.4 |
| 2 | -0.1 | 1.0 | -0.1 | 1.0 | -2.6 | 1.9 | 28.2 | 53.2 | 8.1 | 1.5 |
| 3 | 0.0 | 0.2 | -0.1 | 0.2 | -2.1 | 0.6 | 21.5 | 79.9 | 6.2 | 1.7 |
| 4 | 0.2 | 0.5 | -0.2 | 0.5 | -2.3 | 1.1 | 9.1 | 77.6 | 7.4 | 1.9 |
| 5 | 0.2 | 0.8 | 0.2 | 0.7 | -1.1 | 1.0 | -14.5 | 42.5 | 11.3 | 8.6 |
| 6 | 0.3 | 1.2 | -0.6 | 0.9 | -1.1 | 0.9 | -5.5 | 23.4 | 10.1 | 3.6 |
| 7 | -0.1 | 1.0 | -0.4 | 1.0 | -1.8 | 1.6 | -3.5 | 18.1 | 9.6 | 3.0 |
| 8 | 1.6 | 6.9 | 1.2 | 5.4 | -3.2 | 2.8 | -7.2 | 38.6 | 8.5 | 3.2 |
| 9 | 0.2 | 0.4 | 0.1 | 0.7 | -4.3 | 3.7 | -0.2 | 10.2 | 7.4 | 4.8 |
| 9* | 1.5 | 0.6 | -4.0 | 4.6 | -7.6 | 3.6 | -1.5 | 3.3 | 3.4 | 0.9 |

To test whether including all areas of the input image in the computation of the spectrum would hinder CTF fitting when dealing with photographic films, we also processed set #5 from the CTF challenge (Marabini et al., 2015), which consisted of 17 digitized films including film labels. Despite very strong artefacts in the spectra, CTFFIND4’s defocus parameter estimates were still within 1% of CTFFIND3’s, suggesting our new version may be generally usable, even in those cases.

### 5.3. Quality of fit

To assist in the assessment of its defocus parameter estimates, CTFFIND4 computes an estimate of the maximum resolution at which the agreement between the fit CTF and the experimental signal oscillations is significant (see section 4.3). We chose the threshold for significance (*CC*_*fit*_ = 0.75) heuristically with the goal that the provided estimate correlate well with our visual impression of the fit.

Testing this metric with the CTF challenge datasets, we found that it generally agreed with our visual estimate of the amplitude of Thon rings above background. For example, it tended to report higher fit resolutions for micrographs with carbon than those without (Table 3: compare sets #3, with carbon, and #4, without carbon).

We have also found that it can help diagnose mis-calibrated runtime parameters. For example, the micrographs of set #9, which were simulated *in silico* with a significant amount of astigmatism, clearly have Thon rings extending beyond Nyquist frequency, but our *CC*_*fit*_ = 0.75 criterion initially indicated a good fit only up to 7.4 Å. When runtime parameters were improved (by removing the restraint on astigmatism, and increasing *N*_*d*_ to 512 to avoid CTF aliasing), the mean fit resolution went up to 3.4 Å, and the reported degree of astigmatism (7.6 %) became more consistent with that reported by Marabini et al. (2015, see row 9* of Table 3). This suggests that the *CC*_*fit*_ may be a valid measure of the goodness of fit of the estimated CTF to experimental micrographs.

## 6. Acknowledgments

Our thanks go to Tim Grant for many helpful discussions and suggestions and for reading the manuscript. We’re also grateful to anonymous reviewer #2 of our initial submission, who had several insightful suggestions, particularly regarding Section 4.2.

## Footnotes

1 in some respects, this is similar to what is achieved by phase unwrapping (see e.g. Vargas et al., 2013)

## References

Bartesaghi, A., Matthies, D., Banerjee, S., Merk, A., Subramaniam, S., 2014. Structure of *β*-galactosidase at 3.2Å resolution obtained by cryo-electron microscopy. Proceedings of the National Academy of Sciences 111, 11709–14. doi:10.1073/pnas.1402809111.

Cheng, Y., Grigorieff, N., Penczek, P., Walz, T., 2015. A Primer to Single-Particle Cryo-Electron Microscopy. Cell 161, 438–449. doi:10.1016/j.cell.2015.03.050.

Danev, R., Buijsse, B., Khoshouei, M., Plitzko, J.M., Baumeister, W., 2014. Volta potential phase plate for in-focus phase contrast transmission electron microscopy. Proceedings of the National Academy of Sciences of the United States of America doi:10.1073/pnas.1418377111.

Fernando, K.V., Fuller, S.D., 2007. Determination of astigmatism in TEM images. J Struct Biol 157, 189–200.

van Heel, M., Gowen, B., Matadeen, R., Orlova, E.V., Finn, R., Pape, T., Cohen, D., Stark, H., Schmidt, R., Schatz, M., Patwardhan, a., 2000. Singleparticle electron cryo-microscopy: towards atomic resolution. Quarterly reviews of biophysics 33, 307–69.

van Heel, M., Harauz, G., Orlova, E.V., Schmidt, R., Schatz, M., 1996. A new generation of the IMAGIC image processing system. Journal of structural biology 116, 17–24. doi:10.1006/jsbi.1996.0004.

Huang, Z., Baldwin, P.R., Mullapudi, S., Penczek, P.a., 2003. Automated determination of parameters describing power spectra of micrograph images in electron microscopy. Journal of Structural Biology 144, 79–94. doi:10.1016/j.jsb.2003.10.011.

Mallick, S.P., Carragher, B., Potter, C.S., Kriegman, D.J., 2005. ACE: automated CTF estimation. Ultramicroscopy 104, 8–29.

Marabini, R., Carragher, B., Chen, S., Chen, J., Cheng, A., Downing, K.H., Frank, J., Grassucci, R.a., Heymann, B.J., Jiang, W., Jonic, S., Liao, H.Y., Ludtke, S.J., Patwari, S., Piotrowski, A.L., Quintana, A., Sorzano, C.O., Stahlberg, H., Vargas, J., Voss, N.R., Chiu, W., Carazo, J.M., 2015. CTF Challenge: Result summary. Journal of Structural Biology doi:10.1016/j.jsb.2015.04.003.

McMullan, G., Vinothkumar, K., Henderson, R., 2015. Thon rings from amorphous ice and implications of beam-induced Brownian motion in single particle electron cryo-microscopy. Ultramicroscopy doi:10.1016/j.ultramic.2015.05.017.

Mindell, J.A., Grigorieff, N., 2003. Accurate determination of local defocus and specimen tilt in electron microscopy. Journal of Structural Biology 142, 334–347. doi:10.1016/S1047-8477(03)00069-8.

Penczek, P.A., Fang, J., Li, X., Cheng, Y., Loerke, J., Spahn, C.M.T., 2014. CTER-rapid estimation of CTF parameters with error assessment. Ultramicroscopy 140, 9–19. doi:10.1016/j.ultramic.2014.01.009.

Vargas, J., Otón, J., Marabini, R., Jonic, S., de la Rosa-Trevín, J.M., Carazo, J.M., Sorzano, C.O.S., 2013. FASTDEF: Fast defocus and astigmatism estimation for high-throughput transmission electron microscopy. Journal of Structural Biology 181, 136–148. doi:10.1016/j.jsb.2012.12.006.

Yonekura, K., Braunfeld, M.B., Maki-Yonekura, S., Agard, D.A., 2006. Electron energy filtering significantly improves amplitude contrast of frozen-hydrated protein at 300kV. J Struct Biol 156, 524–536.

